# Characterization of *Brettanomyces bruxellensis* phenolic acid decarboxylase enzyme expressed in *E. coli*

**DOI:** 10.1101/2024.04.16.586637

**Authors:** Michael Lentz

## Abstract

*Brettanomyces* yeasts are considered spoilage organisms in commercial fermentation settings, but some historical beer styles and several modern specialty beer styles rely on *Brettanomyces* for unique aroma and flavor contributions. The organoleptic properties of beer utilizing intentional *Brettanomyces* fermentations can be unpredictable and inconsistent. This study investigates the contributions of *Brettanomyces* fermentations to volatile phenolic character by expression of phenolic acid decarboxylase enzyme cloned from B. *bruxellensis*. Protein was expressed and isolated from *E. coli* cells. The enzyme has broad temperature and pH ranges, and showed variable activity against the most common potential hydroxycinnamic acid substrates.

## Introduction

Beer wort is a rich environment for growth of microorganisms. Successful fermentation to create a desired finished product requires careful attention to yeast selection, and prevention of contamination by unwanted microbes. In commercial fermentation settings, yeasts of the genus *Brettanomyces* are among the most common and detrimental contaminants, leading to undesirable off-aromas and flavors. In the commercial brewery setting, *Brettanomyces* contamination accounts for poorly quantified but significant loss of revenue globally (Suiker & Wosten, 2022).

While *Brettanomyces* infection may be a leading cause of commercial brewery product spoilage, these yeasts may also contribute desirable characteristics under certain conditions for particular beer styles. These include historical styles such as Berliner Weiss, lambic, and Flanders red and brown ales, but also newer “niche” styles, often utilizing *Brettanomyces* in place of or in addition to *Saccharomyces* in a variety of other traditional styles (Crauwels *et al*., 2015; Steensels *et al*., 2015; Colomer *et al*., 2020).

The *Brettanomyces* genus contains five species: *B. bruxellensis, B*. anomalus, *B. custersianus, B. naardenensis*, and *B. nanus* (Shifferdecker *et al*., 2014). A sixth species, *B. acidodurans*, has been proposed for this genus, but not yet confirmed (Peter *et al*., 2017; Suiker & Wosten, 2022). Original isolates of all species came from commercial fermentation settings, however only *B. bruxellensis* and *B. anomalus* are used in brewing unique styles or isolated routinely as a fermentation contaminant (Crauwels *et al*., 2015; Steensels *et al*., 2015; Colomer *et al*., 2020).

Whether desirable or undesirable, one source of aroma and flavor contribution from *Brettanomyces* fermentation is metabolism of hydroxycinnamic acids (HCAs) into their vinyl and ethyl derivatives (Lentz 2018 and references therein). HCAs are phenolic compounds and are components of plant cell walls. They form crosslinks between major cell wall polymers, and bond to other HCA molecules to create lignin and other polyphenols (Vanholm *et al*., 2010; Iiyama *et al*., 1994; Faulds *et al*., 1999). They also provide antimicrobial activity against invading microorganisms (Faulds et al., 1999; Campos et al., 2009). The most common HCAs include ferulic acid, *p*-coumaric acid, caffeic acid, and sinapic acid.

HCAs are released from malted grains and hops into wort during malting, mashing, and boiling (Vanbeneden *et al*., 2007; Vanbeneden *et al*., 2008). The concentration of free HCAs in wort is highly variable, dependent on release from glycosides by enzymes derived from the plant source, and the processes associated with malting and beer production (Vanbeneden *et al*., 2007). They are typically found at concentrations ranging from 1-5 mg/l in <u>wort</u>, but have aroma and flavor thresholds well above this range, typically 20-50 mg/l (Lentz 2018 and references therein). Most commercial fermentation strains of *Saccharomyces* yeast are “phenolic off-flavor negative” (POF-), meaning they are not able to metabolize HCA compounds. Selection for POF-strains was a major driving force in the domestication of brewing and winemaking strains of yeast (Gallone *et al*., 2016). Many wild strains of yeast, including wild *Saccharomyces* strains and all *B. bruxellensis* and *B. anomalus* isolates tested encode a phenolic acid decarboxylase (PAD) enzyme that converts one or more HCAs to their vinyl derivatives (Clausen *et al*., 1994; Edlin *et al*., 1998; Godoy *et al*., 2008; Cavin *et al*., 1996; Cavin *et al*., 1998). A smaller number of microbes, including *Brettanomyces*, also encode a vinylphenol reductase that converts the vinyl intermediates to their ethyl derivatives (Chatonnet *et al*., 1992; Romano *et al*., 2017). Both the vinyl and ethyl compounds created by these enzymes have very low flavor and aroma thresholds. Most are considered undesirable and are described as medicinal, plastic, rubbery, etc. They may also be described as smoky, spicy, clove-like, or leathery, features that may be desirable at certain levels in specific beer styles (White & Zainasheff, 2010).

Even when used intentionally in a brewery setting, use of *Brettanomyces* strains is unpredictable compared to the well-characterized brewing strains of *Saccharomyces*. To better understand the role of the PAD enzyme in flavor and aroma contributions to beer and wine, we have cloned the PAD gene from a commercially available brewing strain. Here we describe properties of the enzyme isolated from *E. coli* cells overexpressing the cloned PAD gene.

## Materials and Methods

### Cloning

PCR primers were designed using PAD gene sequences determined from published whole-genome sequences (genbank accession numbers GCA_000688595.1, GCA_000340765.1, GCA_000259595.1). Primers included restriction enzyme sites to facilitate cloning (Table 1). Primers were synthesized by Invitrogen.

**Table 1.**
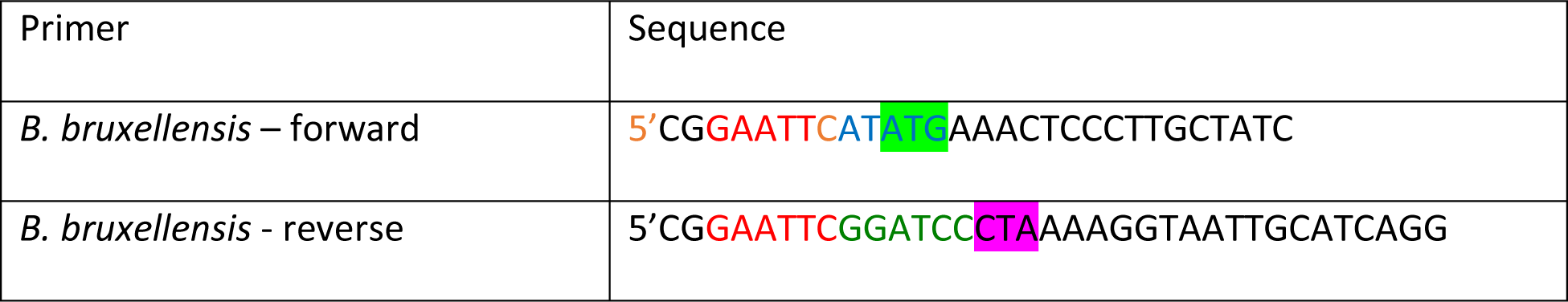
Primer sequences used for cloning. Colored letters represent restriction enzyme sites and highlights represent start (forward) and stop (reverse) codons.

50 ul PCR reactions were assembled that included 0.5 uM of each primer and 1X PCR MasterMix (Thermo Scientific). For DNA template, 0.5 ul of overnight liquid culture of yeast cells was added directly to the PCR reaction. Reaction protocol used was:

1. 95°C, 10 min
2. 95°C 1 min
3. 45°C 1 min
4. 72°C 2 min
5. Go to step 2, X34 6.
6. 72°C 10 min

PCR was confirmed by gel electrophoresis. 5 ul PCR product was digested with BamH1 and Nde1 restriction enzymes according to manufacturer directions. PET vector DNA was also digested as for the PCR products, followed by treatment with shrimp alkaline phosphatase according to the manufacturer instructions. Digestion products were isolated by gel electrophoresis and purified using a GeneClean kit protocol (MP Biomedicals). Ligation of the PAD gene into the pET vector was accomplished by standard protocols (Sambrook and Russell, 2001) and transformed into *E. coli* DH5alpha cells according to supplier directions (Thermo Scientific). DNA was isolated from putative clones and confirmed by restriction enzyme digestion and electrophoresis. All cloning enzymes were from Thermo Scientific.

### PAD protein overexpression

pET-PAD plasmids were transformed into competent *E. coli* BL21DE3 cells according to the manufacturer (Thermo Scientific). Single colonies were isolated and used to inoculate 5 ml overnight cultures in LB broth with ampicillin. 1 ml of stationary phase overnight culture was inoculated into 100 ml LB broth with ampicillin and grown at 37°C for 3 hours. Isopropyl-beta-D-thiogalactopyranoside (IPTG) was added to 1 mM and the temperature reduced to 25°C for growth overnight. Cells were harvested by centrifugation at 5000 rpm for 10 min at 4°C. Pellets were washed in one-half volume of cold phosphate-buffered saline (PBS) and centrifuged as before. Cells were resuspended in 10 ml cold PBS followed by addition of phenymethylsulfonyl flouride to 1 nM. Cells were broken by 4 × 20 sec, 12 W sonication cycles using a probe sonicator (Misonix). Extracts were clarified by centrifugation at 10,000 rpm for 30 min. Protein extracts were stored at 4°C or -20C, or diluted to 50% in sterile glycerol and stored at -20°C.

### Enzyme assay

Reactions were prepared in 1.5 ml microcentrifuge tubes. 10 ul protein extract and 10 ul 20 mM HCA were added to 80 ul phosphate-buffered saline (PBS) at pH 6.0 and mixed briefly. Reactions were incubated at room temperature for 60 min. 20 ul reaction mix was added to 180 ul H_2_O in a 96 well, flat-bottom, UV transparent microplate. Sample controls included water (blank) and 0.2 mM HCA (negative control). Samples were read at 287 nm for *p*-coumaric acid and sinapic acid, or 313 nm for ferulic acid and caffeic acid (Cavin *et al*., 1997). These wavelengths correspond to the peak absorbance values of the substrates (data not shown). Reduction of the absorbance value corresponds to metabolism of the HCA to vinyl product. For analysis of PAD enzyme pH tolerance, the PBS is the reaction was replaced with acetate, phosphate, or Tris buffer at the indicated pH.

## Results and Discussion

### Cloning

The PAD gene was identified from several whole genome sequencing projects for several strains of *B. bruxellensis* (genbank accession numbers GCA_000688595.1, GCA_000340765.1, GCA_000259595.1). It was noted that not all strains have the start codon identified by Godoy *et al*. (2014) in their cloned PAD gene. We therefore used the start site of the “PAD2” sequence later identified by Gonzalez *et al*. (2017). The sequences analyzed varied by zero to three nucleotides across the gene sequence. The 5’ and 3’ ends of the coding sequence were therefore used to design PCR primers for amplification of the gene from a *B. bruxellensis* commercial brewing strain. Using yeast cells directly from liquid culture as a source of template DNA yielded the most consistent PCR reaction products. PCR products were subject to restriction enzyme digestion followed by ligation into linearized pET3a expression vector (pET-PAD). Clones were screened by gel electrophoresis and restriction enzyme analysis.

### *E. coli* growth/expression

*E. coli* BL21DE3 cells were transformed with the pET-PAD clone or empty vector as a control. Separate cultures of PAD or negative control cells were induced for PAD expression and “challenged” with 2 mM of one of four HCAs, and growth was monitored for 28 hours. Interestingly, cells expressing PAD responded differently to the various HCAs, unlike the control cells which grew at the same rate regardless of HCA addition. As shown in figure 1, PAD expressing cells showed a slowed growth curve in the presence of p-coumaric acid and ferulic acid compared to sinapic acid, which did not slow growth. The growth rate in the presence of caffeic acid was intermediate.

**Figure 1.**
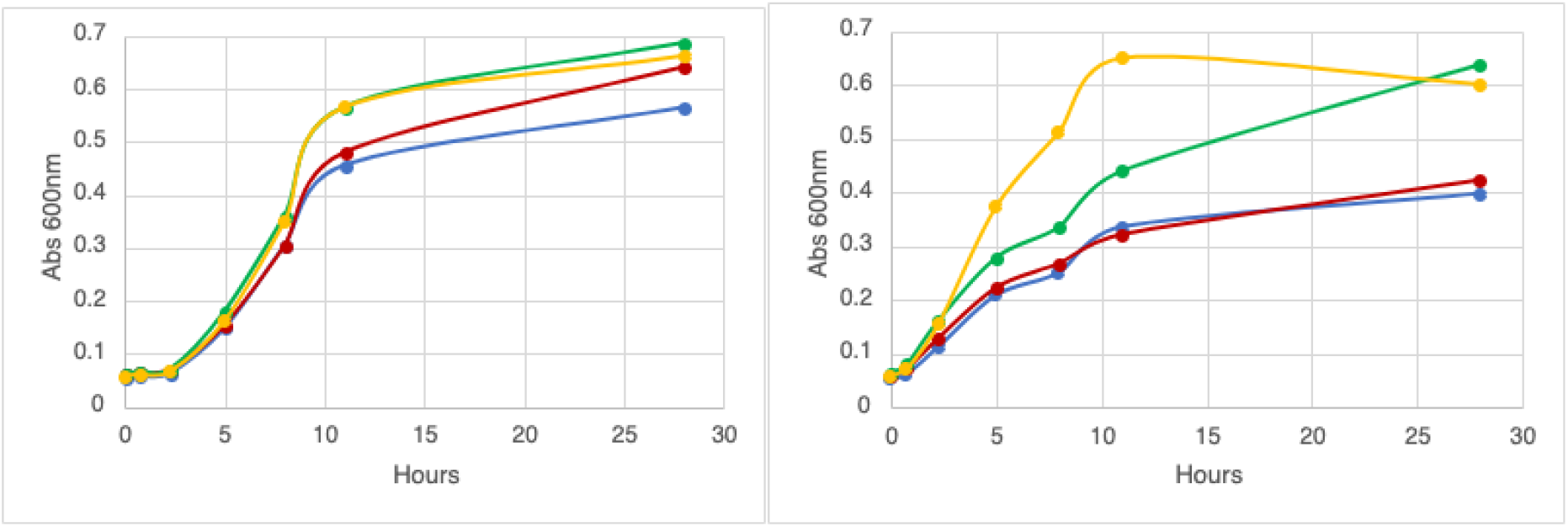
*E. coli* cells expression empty vector (left) or *B. bruxellensis* PAD enzyme (right) were monitored for growth rate over 28 hours in one of four HCAs (blue, ferulic acid; red, *p*-coumaric acid, green, caffeic acid, orange, sinapic acid).

It was unexpected that *E. coli* cells expressing PAD enzyme were growth inhibited in substrate that they should be able to metabolize, compared to control cells that were unable to metabolize the HCAs but were not growth inhibited. It might be expected that growth inhibition would be higher in cells that were unable to metabolize the toxic HCA. One possible explanation is that the energy required for high level expression of a single enzyme reduces the overall metabolic fitness of the cells, accounting for the observed slower growth, however this explanation does not account for growth differences in the presence of different HCAs.

Samples of culture medium from the growth experiment were also analyzed for metabolism of the HCAs present. The data shown in figure 2 indicates that *E. coli* cells were converted from a PAD-phenotype to PAD+ by induction of the PAD gene vector. Cells carrying the empty vector were unable to convert any of the four substrates tested to the corresponding vinyl derivatives, as measured by monitoring absorption at wavelengths corresponding to the peak value for each substrate. PAD expressing cells metabolize three of the four HCAs, as measured by reduction in absorbance by the substrate molecules. Both *p*-coumaric acid and ferulic acid were readily metabolized, while caffeic acid was converted at a slower rate. The PAD expressing cells were unable to metabolize sinapic acid.

**Figure 2.**
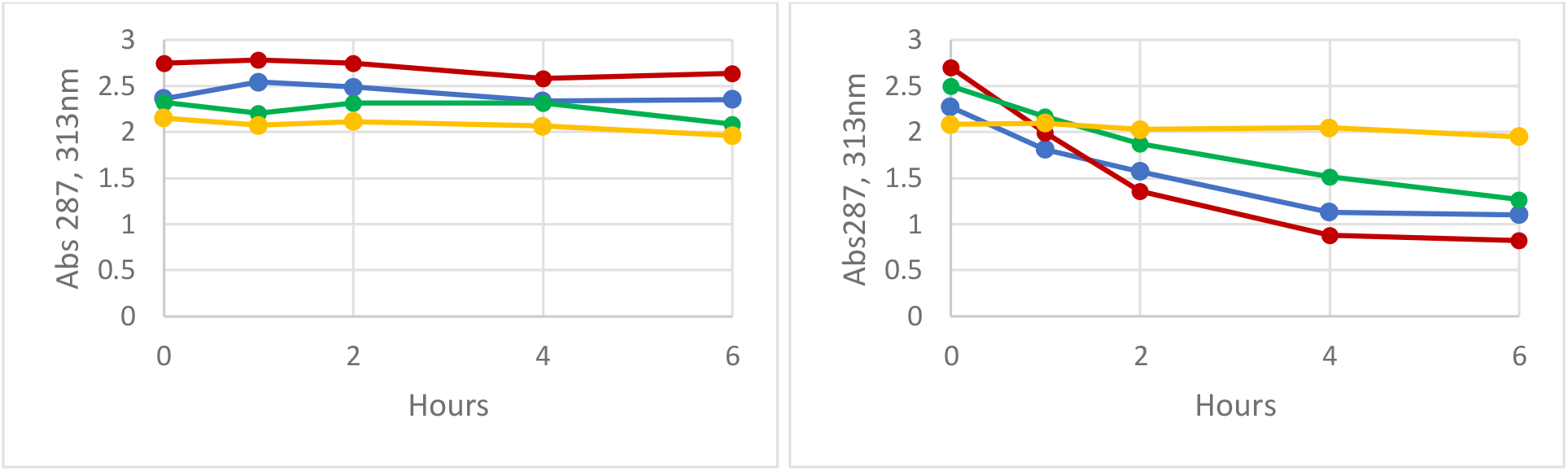
*E. coli* cells expressing PAD enzyme (right) or harboring empty vector (left) were grown in each of four HCAs (blue, ferulic acid; red, *p*-coumaric acid, green, caffeic acid, orange, sinapic acid). PAD activity was monitored over six hours by measuring absorbance at 313nm. Reduced absorbance indicates metabolism of the substrate.

### PAD protein activity

Cultures of PAD expressing *E. coli* cells or empty vector controls were induced and allowed to express protein for 18 hours. Protein extracts were then prepared from harvested cells. Extracts were incubated with each of four HCA substrates and monitored for metabolism over two hours. Similar to metabolism by live *E. coli* cells, extracts readily converted ferulic acid and *p*-coumaric acid to their vinyl products. Caffeic acid was converted to an intermediate extent, and sinapic acid was not metabolized. None of the substrates was metabolized by the empty vector negative control protein extract (figure 3).

**Figure 3.**
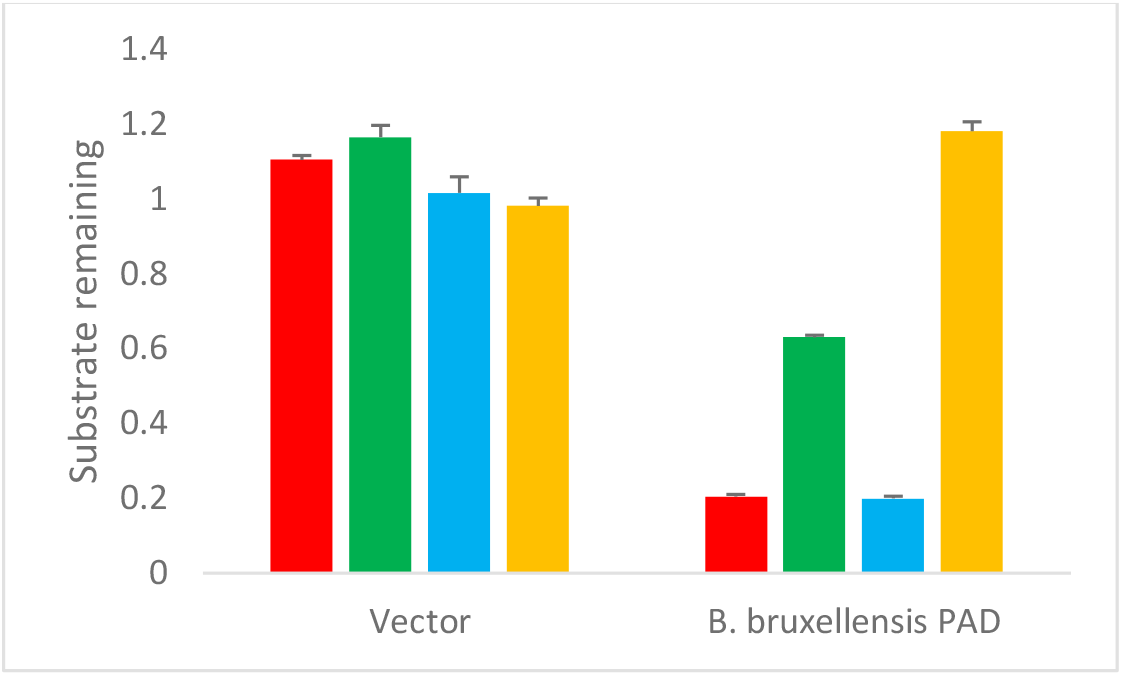
PAD activity of protein extracts. Extracts prepared from either empty vector cells (left) or PAD gene expressing cells were incubated with each of four potential substrates (blue, ferulic acid; red, *p*-coumaric acid, green, caffeic acid, orange, sinapic acid). Data represents substrate remaining after one hour of incubation. Error bars, standard error of the mean of three trials.

PAD protein extracts were incubated with ferulic acid in buffered solutions ranging from pH 4 to pH 8.8 and monitored for activity as described above. The enzyme has a broad pH tolerance, with significant activity from pH 4.8 to 8.0. Maximum activity was observed between pH 5.0 and 7.0. The results are in general agreement with data published on PAD enzyme purified directly from *B. bruxellensis* yeast cells (Godoy *et al*., 2008). The temperature optimum for the enzyme was 40°C and was active over a range of 10C through 50°C.

## Conclusions

Phenolic acid decarboxylase genes from several microbial species have been cloned and protein expressed in heterologous systems. The *Bacillus subtilis* PAD was cloned and overexpressed in *E. coli* to investigate enzyme features related to substrate specificity (Cavin *et al*., 1998). The PAD enzyme from the Koji mold *Aspergillus luchuensis* was likewise cloned and expressed in *E. coli* from the pET vector system to investigate properties of the enzyme and its possible role in the organoleptic qualities of fermented awamori beverage (Maeda *et al*., 2018). The *B. bruxellensis* PAD gene was previously cloned and expressed in a POF-*S. cerevisiae* strain to confirm cloning (Godoy *et al*., 2014), and more recently to investigate differences the reaction mechanisms between the *B. bruxellensis* and *S. cerevisiae* PAD enzymes (Ogata and Saito, 2024).

We have adapted the widely used pET bacterial expression system for production and analysis of *Brettanomyces* phenolic acid decarboxylase enzyme in *E. coli* to investigate this enzyme’s role in phenolic flavor and aroma production in beer and wine fermentations.

The variation observed in metabolism of the four tested HCAs can partially be explained by the structure of the substrate molecules. All four share the cinnamic acid structure (acrylic acid bound to a phenol ring). The four HCAs differ in additional substitutions at other positions on the ring. *p*-coumaric acid has a single hydroxyl group opposite the acrylic group and is readily metabolized. Sinapic acid, which does not function as substate, contains two bulky methoxy groups in addition to the hydroxyl group at the meta positions. These additional groups may prevent proper entry or binding of this compound to the active site of the enzyme. Ferulic acid, which is metabolized as efficiently as p-coumaric acid, has one of the two methoxy groups found on sinapic acid, but not the second. Interestingly, caffeic acid, a relatively poor substrate compared to ferulic and p-coumaric acids, has a smaller hydroxyl group at the position occupied by the methoxy group in ferulic acid. Therefore it cannot be simply a matter of “fit” that determines the suitability of an HCA as a substrate for this enzyme. Structural analysis and substrate modeling may help to answer this question.

This bacterial expression system can now be used to further investigate the properties of *Brettanomyces* PAD enzyme and its role in volatile phenol production in commercial fermentations.

## Acknowledgements

The author acknowledges Y. Barns and W. Smith for some of the data collection. This work was supported by the American Society for Brewing Chemists Research Council.

The author reports there are no competing interests to declare.

## References

Cavin J-F, Barthelmebs L, Guzzo J, Van Beeumen J, Samyn B, Travers J-F, Divies C. (1997) Purification and characterization of an inducible p-coumaric acid decarboxylase from Lactobacillus plantarum. FEMS Microbiology Letters 147; 291–295. 10.1111/j.1574-6968.1997.tb10256.x

Cavin J-F, Dartois V, Divies C. (1998) Gene cloning, transcriptional analysis, purification, and characterization of phenolic acid decarboxylase from Bacillus subtilis. Applied and Environmental Microbiology 64; 1466–1471. DOI: 10.1128/AEM.64.4.1466-1471.1998

Chatonnet P, Dubourdie D, Boidron J, Pons M. (1992) The origin of ethylphenols in wines. Journal of the Science of Food and Agriculture 60 (2); 165–178 10.1002/jsfa.2740600205

Clausen M, Lamb CJ, Megnet R, Doerner PW. (1994) PAD1 encodes phenylacrylic acid decarboxylase which confers resistance to cinnamic acid in Saccharomyces cerevisiae. Gene 142; 107–112. 10.1016/0378-1119(94)90363-8

Colomer MS, Chailyan A, Fennessy RT, Olsson KF, Johnsen L, Solodovnikova N and Forster J (2020) Assessing population diversity of Brettanomyces yeast species and identification of strains for brewing applications. Frontiers in Microbiology 11:637. doi: 10.3389/fmicb.2020.00637

Crauwels S, Steensels J, Aerts G, Willems KA, Verstrepen KJ, Lievens B. (2015) Brettanomyces bruxellensis, essential contributor in spontaneous beer fermentation providing novel opportunities for the brewing industry. Brewing Science 68; 110–121.

Degrassi G, Polverino de Laureto P, Bruschi CV. (1995) Purification and characterization of ferulate and p-coumarate decarboxylase from Bacillus pumilus. Applied and Environmental Microbiology 61; 326–332. DOI: 10.1128/aem.61.1.326-332.1995

Gallone B, Steensels J, Prahl T, Soriaga L, Saels V, Herrera-Malaver B, Merlevede A, Roncoroni M, Voordeckers K, Miraglia L, Teiling C, Steffy B, Taylor M, Schwartz A, Richardson T, White C, Baele G, Maere S, Verstrepen KJ. (2016) Domestication and divergence of Saccharomyces cerevisiae beer yeasts. Cell 166 (6); 1397-1410. e16, ISSN 0092-8674, 10.1016/j.cell.2016.08.020.

Godoy L, Martínez C, Carrasco N, Ganga MA. (2008) Purification and characterization of a p-coumarate decarboxylase and a vinylphenol reductase from Brettanomyces bruxellensis. International Journal of Food Microbiology 127; 6–11. 10.1016/j.ijfoodmicro.2008.05.011

Godoy L, García V, Peña R, Martínez C, Ganga MA. (2014) Identification of the Dekkera bruxellensis phenolic acid decarboxylase (PAD) gene responsible for wine spoilage. Food Control 45; 81–86. ISSN 0956-7135, 10.1016/j.foodcont.2014.03.041.

Gonzalez C, Godoy L, Ganga MA. (2016) Identification of. Second PAD1 in Brettanomyces bruxellensis LAMAP2480. Antonie van Leeuwenhoek 110; 291–296 DOI 10.1007/s10482-016-0793-3

Huang H-K, Tokashiki M, Maeno S, Onaga S, Taira T, Ito S. (2012) Purification and properties of phenolic acid decarboxylase from Candida guilliermondii. Journal of Industrial Microbiology and Biotechnology 39; 55–62. 10.1007/s10295-011-0998-4

Lentz M. (2018) The impact of simple phenolic compounds on beer aroma and flavor. Fermentation 4 (20). doi:10.3390/fermentation4010020

Maeda M, Tokashiki M, Tokashiki M, Uechi K, Ito S, Taira T. (2018) Characterization and induction of phenolic acid decarboxylase from Aspergillus luchuensis. Journal of Bioscience and Bioengineering 126 (2); 162–168. 10.1016/j.jbiosc.2018.02.009

Ogata T, Saito M. (2024) Differences in formation mechanisms of phenolic off-flavor compounds among yeast species. Journal of the American Society of Brewing Chemists, 82:1, 61–65. DOI: 10.1080/03610470.2023.2193921

Parada-Fabian JC, Hernandez-Sanchez H, Mendez-Tenorio A. (2019) Substrate specificity of the phenolic acid decarboxylase from Lactobacillus plantarum and related bacteria analyzed by molecular dynamics and docking. Journal of Plant Biochemistry and Biotechnology 28 (1); 91–104. 10.1007/s13562-018-0466-6

Péter G, Dlauchy D, Tobias A, Fülop L, Podgorsek M, Cadez N. (2017) Brettanomyces acidodurans sp. Nov., a new acetic acid producing yeast species from olive oil. Antonie van Leeuwenhoek 110; 657–664 doi:10.1007/s10482-017-0832-8

Rodriguez H, Angulo I, de las Rivas B, Campillo N, Paez JA, Muñoz R, Mancheño JM. (2010) p-Coumaric acid decarboxylase from Lactobacillus plantarum: Structural insights into the active site and decarboxylation catalytic mechanism. Proteins 78; 1662–1676. DOI: 10.1002/prot.22684

Romano D, Valdetara F, Zambelli P, Galafassi S, De Vitis V, Molinari F, Compagno C, Foschino R, Vigentini I. (2017) Cloning the putative gene of vinyl phenol reductase of Dekkera bruxellensis in Saccharomyces cerevisiae. Food Microbiol. 63;92–100. doi: 10.1016/j.fm.2016.11.003. Epub 2016 Nov 3. PMID: 28040186.

Schifferdecker JA, Dashko S, Ishchuk OP, Piskur J. (2014) The wine and beer yeast Dekkera bruxellensis. Yeast 31(9); 323–332. DOI: 10.1002/yea.3023

Sheng X, Lind MES, Himo F. (2015) Theoretical study of the reaction mechanism of phenolic acid decarboxylase. FEBS Journal 282; 4703–4713. 10.1111/febs.13525

Suiker IM, Wosten HAB. (2022) Spoilage yeasts in beer and beer products. Current Opinion in Food Science 2022, 44:100815 10.1016/j.cofs.2022.100815

